# A framework for disentangling ecological mechanisms underlying the island species-area relationship

**DOI:** 10.1101/410126

**Authors:** Jonathan M. Chase, Leana Gooriah, Felix May, Wade A. Ryberg, Matthew S. Schuler, Dylan Craven, Tiffany M. Knight

## Abstract

The relationship between an island’s size and the number of species on that island—the island species-area relationship (ISAR)—is one of the most well-known patterns in biogeography, and forms the basis for understanding biodiversity loss in response to habitat loss and fragmentation. Nevertheless, there is contention about exactly how to estimate the ISAR, and the influence of the three primary ecological mechanisms—random sampling, disproportionate effects, and heterogeneity— that drive it. Key to this contention is that estimates of the ISAR are often confounded by sampling and estimates of measures (i.e., island-level species richness) that are not diagnostic of potential mechanisms. Here, we advocate a sampling-explicit approach for disentangling the possible ecological mechanisms underlying the ISAR using parameters derived from individual-based rarefaction curves estimated across spatial scales. If the parameters derived from rarefaction curves at each spatial scale show no relationship with island area, we cannot reject the hypothesis that ISARs result only from random sampling. However, if the derived metrics change with island area, we can reject random sampling as the only operating mechanism, and infer that effects beyond sampling (i.e., disproportionate effects and/or heterogeneity) are also operating. Finally, if parameters indicative of within-island spatial variation in species composition (i.e., β-diversity) increase with island area, we can conclude that intra-island compositional heterogeneity plays a role in driving the ISAR. We illustrate this approach using representative case studies, including oceanic islands, natural island-like patches, and habitat fragments from formerly continuous habitat, illustrating several combinations of underlying mechanisms. This approach will offer insight into the role of sampling and other processes that underpin the ISAR, providing a more complete understanding of how, and some indication of why, patterns of biodiversity respond to gradients in island area.

## Introduction

The relationship between the area sampled and the number of species in that area—the species-area relationship—is one of the oldest laws in ecology (e.g., Arrhenius 1921, Lawton 1999, Lomolino 2000, Drakare et al. 2006). There are many forms of SARs that represent rather distinct patterns and processes (e.g., Scheiner 2003, Scheiner et al. 2011), but we here focus specifically on one type, the Island Species-Area Relationship (hereafter ISAR). The ISAR correlates how the numbers of species (species richness) varies with the size of islands or distinct habitat patches (natural or fragmented due to human activities). Like other types of SARs, the ISAR is usually positive (e.g., MacArthur and Wilson 1963, 1967, Connor and McCoy 1979, Triantis et al. 2012). However, complexities such as island age, habitat heterogeneity and/or isolation can complicate this simple expectation (Kreft et al. 2008, Borregard et al. 2016).

We refer to ‘islands’ in the ISAR as any insular system, including true islands, habitat patches that are surrounded by distinctly different habitats (matrix) (e.g., lakes, edaphically delimited habitats), and habitat fragments that have been insularized by human activities. In addition to being an important biogeographic pattern in its own right, the ISAR and concepts closely related to it play an important role in understanding how biodiversity changes when habitat is lost and/or fragmented into smaller island-like habitats (e.g., Diamond 1975, Simberloff and Abele 1976, Matthews et al. 2014, 2016, Fahrig 2017). As a result, understanding the patterns and the processes underlying ISARs and their derivatives would seem to be an important endeavor in the context of island biogeography and conservation.

Despite its conceptual importance, there remains a great deal of ambiguity regarding ISAR patterns, as well as its underlying processes (e.g., Scheiner et al. 2011). When describing ISAR patterns, authors report and analyze different aspects of species richness regressed against total island size, including total numbers of species and the number of species found within a constantly-sized sub-sampled area. Such different sampling designs have created confusion when comparing slopes of ISARs; an increasing number of species measured in a fixed-area plot with increasing island area means something quite different than an increasing number of species on the entire island (see also Hill et al. 1994, Gilaldi et al. 2011, 2014). In terms of processes underlying the ISAR, there is similar confusion. Multiple mechanisms, including passive sampling, colonization/extinction (i.e., metacommunity) dynamics, and habitat heterogeneity, as well as their interactions, have been invoked to explain ISARs. Unfortunately, the exact ways by which these mechanisms operate, and how they can be disentangled from observational data, remains in question.

Following others (e.g., Triantis et al. 2012, Mathews et al. 2014, 2016), we refer to the ISAR as the relationship between the total species richness on a given island (or habitat patch) and the size of that island. However, simply knowing the shape of the relationship between the size of an island and the total species richness (hereafter S_total_) on that island can tell us very little about the possible mechanisms underlying the ISAR. In order to understand the mechanisms underlying the ISAR, it is necessary to collect and analyze data at the level below the scale of the entire island (see also Hill et al. 1994, Yaacobi et al. 2007, Stiles and Scheiner 2010, Gilaldi et al. 2011, 2014). Specifically, we recommend collecting data from multiple standardized plots where both the numbers and relative abundances of species are available, as well as compositional differences of species among locations within an island. We recognize that this requires extra data often not available for many biogeographical and macroecological studies of island systems, but emphasize that the extra effort involved allows a much deeper understanding of the possible processes underlying the ISAR patterns observed.

## Mechanisms Underlying the ISAR

We overview three general classes of potential mechanisms underlying the ISAR—passive sampling, disproportionate responses and heterogeneity—from least complex to most complex (see also Connor and McCoy 1979, McGuinness 1984, Scheiner et al. 2011 for deeper discussions of these mechanisms more generally for all types of SARs). Then we discuss how they can be detected using a multi-scale, multi-metric approach. Importantly, there remains much confusion in the literature regarding exactly which mechanisms can create the ISAR, which patterns these mechanisms generate, and how to disentangle them. Thus, we begin with a general overview of the general classes of mechanisms and discuss how they can be disentangled with a more directed sampling approach.

In brief, ***passive sampling*** (sometimes called the ‘more individuals hypothesis’) emerges when larger islands have more species than smaller islands via passive sampling of individuals (and thus species) from a larger regional pool. ***Disproportionate response*** (sometimes called ‘area per se’) include a large array of possible mechanisms that influence the likelihood that some species are favored, and others disfavored, on islands of different sizes, such that they achieve different relative abundances on different sized islands. ***Heterogeneity*** also leads to disproportionate responses and altered relative abundances of species, but these emerge at larger scales via clumping of species that can emerge because of habitat differences and/or dispersal limitation. In the following sections, we discuss each of these mechanisms, and possible ways to detect them from within-island surveys.

### Passive sampling

The simplest mechanism of the ISAR is that islands passively sample individuals from a larger ‘regional’ pool of individuals of different species. Larger islands passively sample more individuals, and thus more species, from the regional pool. This is essentially a ‘null’ hypothesis, but one that can be tested using standard methods, and which provides important insights about the potential underlying processes leading the ISAR. The influence of passive sampling on the ISAR was first described by Arrhenius (1921) in one of the first quantitative explorations of the ISAR. It is important to emphasize that sampling effects are sometimes thought of as an artifact of limited sampling for uncovering the true numbers of species. This is not the case for this passive sampling null hypothesis. It is also implicit in several early quantitative explorations of the ISAR where the regional pool consists of few common and many rare species, and smaller islands passively sample fewer individuals, and thus fewer species than larger islands (i.e., Preston 1960, May 1975).

Coleman (1981) developed an analytical formula for this process based on random placement of individuals on islands and Coleman et al. (1982) applied it to data from samples of breeding birds on islands in a reservoir to suggest that this passive sampling mechanism most likely explained the ISAR in this system. This will create a positive ISAR with more rare species being present on larger islands, but only in proportion to their abundance in the total pool (i.e., the relative proportions of species does not change from small to large islands). Importantly, this random placement method is nearly identical to individual-based rarefaction methods (e.g., Gotelli and Colwell 2001), which we use below to test the random sampling hypothesis.

Several authors have tested the passive sampling hypothesis by measuring the numbers of species in a given fixed area on islands of different sizes and correlating that density with the total area of the island (e.g., Hill et al. 1994, Kohn and Walsh 1994, Yaacobi et al. 2007, Gilaldi et al. 2011, 2014). If the number of species in a fixed area sample does not vary as island size varies, this is taken to imply that passive sampling is most likely the only mechanism acting. However, if the number of species in a fixed area increases as island size increases, we would instead conclude that there is some biological effect, beyond sampling, that allows more species to persist in a given area on larger than smaller islands.

While fixed-area sampling can be useful for inferring whether ISAR patterns deviate from patterns expected from pure sampling effects, this method is unfortunately not as powerful of a ‘null hypothesis’ as has often been suggested. There are at least two common factors that can lead to patterns that appear consistent with the passive sampling hypothesis that in fact emerge from effects that are beyond sampling. First, when disproportionate effects are primarily experienced by rare species, sampling at small spatial grains may miss this effect, especially when averages of the numbers of species are taken from the smallest spatial scale. For example, Karger et al. (2014) found that fern species richness in standardized plots did not increase with island area when measured at small spatial grains (i.e., 400m^2^-2400m^2^), but that the slope significantly increasing at the largest sampling grain (6400 m^2^). Second, it is possible that species richness measured in standardized plots may not vary with island size, but that habitat heterogeneity leads to different species present in different habitat types, creating the ISAR. For example, Sfenthourakis and Panitsa (2012) found that plant species richness on Greek islands measured at local (100m^2^) scales did not change with island area, but that there were high levels of β-diversity on islands that were larger, likely due to increased heterogeneity. In both of these studies, simply measuring standardized species richness in small plots across islands of different spatial grains may have led to the faulty conclusion of random sampling effects.

### Disproportionate effects

When disproportionate effects underlie the ISAR, there are more species on larger islands because species from the regional pool are differentially influenced by island size (as opposed to the passive sampling hypothesis, where species are proportionately influenced by island size). Disproportionate effects includes a number of different sub-mechanisms whereby some species are favored, and others disfavored, by changes in island size.

Most such mechanisms predict that the numbers of species in a fixed sampling area should increase with increasing island size (sometimes called ‘area per se’ mechanisms; Connor and McCoy 1979). The mostly widely considered of these mechanisms is MacArthur and Wilson’s (1963, 1967) theory of island biogeography. Here, the rates of colonization of species increases with island size, and the rates of extinction decrease with island size, leading to the expectation that more species should often be able to persist in a fixed area on larger islands. Several other kinds of spatial models can also predict similar patterns whereby the coexistence of several species is favored when the total area increases (e.g., Hanski et al. 2013), or when population-level processes, such as Allee-effects or demographic stochasticity, are less likely on larger relative to smaller islands (e.g., Hanski and Gyllenberg 1993, Orrock and Wattling 2010). Disproportionate effects can also emerge when changes in island size influences island-level environmental conditions. For example, smaller islands are often more likely to experience disturbances and/or have lower productivity (McGuinness 1984), and in the context of habitat fragmentation, smaller island fragments often have edge effects whereby habitat-specialist species are negatively impacted (Ewers and Didham 2006).

Although often less well appreciated, mechanisms similar to those described above can favor multiple species in smaller, rather than larger habitats. For example, it is possible that more widespread species can dominate larger habitats via high rates of dispersal and mass effects. Likewise, especially in the context of habitat islands formed via habitat fragmentation, disproportionate effects favoring species in smaller islands can include the disruption of exclusion interspecific interactors (e.g., pathogens, predators or competitors), or more species favored by edges and heterogeneity created in smaller habitats (Fahrig 2017). In such cases, we might expect a weaker or even negative ISAR depending on whether random sampling effects (which are always operating) outweigh the disproportionate effects

### Heterogeneity

The last family of mechanisms that have been proposed to lead to the ISAR involve heterogeneity in the composition of species within islands. These mechanisms are centered on the supposition that larger islands can have more opportunity for species to aggregate intraspecifically or clump (leading to heterogeneity in species composition) than smaller islands. This can emerge from two distinct sub-mechanisms:

(i) *Habitat heterogeneity*. Habitat heterogeneity leads to dissimilarities in species composition via the ‘species sorting’ process inherent to niche theory (e.g., Whittaker 1970, Tilman 1982, Chase and Leibold 2003). As a mechanism for the ISAR, larger islands are often assumed to have higher levels of habitat heterogeneity than smaller islands (e.g., Williams 1964, Hortal et al. 2009). For example, larger oceanic islands typically have multiple habitat types, including mountains, valleys, rivers, etcetera, allowing for multiple types of species to specialize on these habitats, whereas smaller islands only have a few habitat types. Likewise, in freshwater lakes, which can be thought of as aquatic islands in a terrestrial ‘sea’, larger lakes typically have more habitat heterogeneity (e.g., depth zonation) than smaller lakes.
(ii) *Compositional heterogeneity due to dispersal limitation*. Dispersal limitation can also lead to compositional heterogeneity through a variety of spatial mechanisms, including ecological drift, colonization and competition tradeoffs, and the like (e.g., Condit et al. 2002, Leibold and Chase 2017). If dispersal limitation is more likely on larger islands, we might expect greater within-island spatial coexistence via dispersal limitation, higher compositional heterogeneity, and thus greater total species richness on larger than on smaller islands.

Patterns of species compositional heterogeneity that emerge from these two distinct mechanisms are difficult to distinguish without explicit information on the characteristics of habitat heterogeneity itself, as well as how species respond to that heterogeneity. While we do not explicitly consider it further here, the spatial versus environmental drivers of compositional heterogeneity (β-diversity) can be more acutely disentangled if site-level environmental conditions and spatial coordinates are known by using standard methods in metacommunity ecology (e.g., Peres-Neto et al. 2006, Ovaskanien et al. 2017).

Finally, as with disproportionate effects above, opposite patterns are also possible. While we typically assume that heterogeneity increases with island area, leading to the positive ISAR, this need not be true. For example, smaller islands have higher perimeter:area ratios (i.e., edge effects), and thus can have higher levels of heterogeneity than larger islands by some measures.

## Disentangling ISAR Mechanisms with Observational Data

As a result of the often impracticality of field experiments on the ISAR at realistic scales (but see Simberloff 1976), considerable attention has been paid towards developing sampling and analytical methodology that can allow a deeper understanding of potential ISAR mechanisms from observational data. However, these approaches have appeared piecemeal in the literature, are incomplete, and have not yet been synthesized into a single analytical framework.

Furthermore, two or more of these mechanisms can act in concert and are non-exclusive (e.g‥, Chisholm et al. 2016). For example, the influence of passive sampling is likely always occurring in the background even when disproportionate effects and/or heterogeneity also influence ISAR patterns. Thus, even if we reject passive sampling as the sole mechanism leading to the ISAR via deviations from the null expectation, we cannot say that passive sampling does not at least partially influence the observed patterns. The same is true for any null modelling approach. Likewise, it is possible that disproportionate responses of species via alterations to spatial or local conditions can act in concert with changes in habitat heterogeneity. In this case, however, we can more completely falsify these processes by comparing patterns both within communities (α-diversity) and among communities (β-diversity), as we will discuss in more detail below.

Here, we overview a generalized approach for disentangling the possible mechanisms underlying the ISAR. Our approach is based on recent work that uses an individual-based rarefaction framework (e.g., Gotelli and Colwell 2001) to calculate several measures of biodiversity at multiple spatial scales (e.g., Chase et al. 2018, McGlinn et al. 2018). And then to relate these measures to variation in island size. In a sense, then, we propose the use of within-island species richness relationships (Type II or Type III curves from Scheiner 2003, Scheiner et al. 2011) to evaluate the mechanisms underlying among-island ISAR relationships (Type IV curves from Scheiner 2003, Scheiner et al. 2011).

Figure 1a overviews the sampling design necessary on an island in order to calculate the parameters necessary to disentangle ISAR mechanisms. Specifically, in addition to estimating the total numbers of species on an island (S_total_), we advocate sampling multiple standardized plots within a given island (ideally stratified across the island and any potential habitat heterogeneity) so that a number of parameters can be derived and compared with island size. These parameters are described in Table 1 and can be visualized as components along individual-based rarefaction curves as in Figure 1b.

**Table 1.**
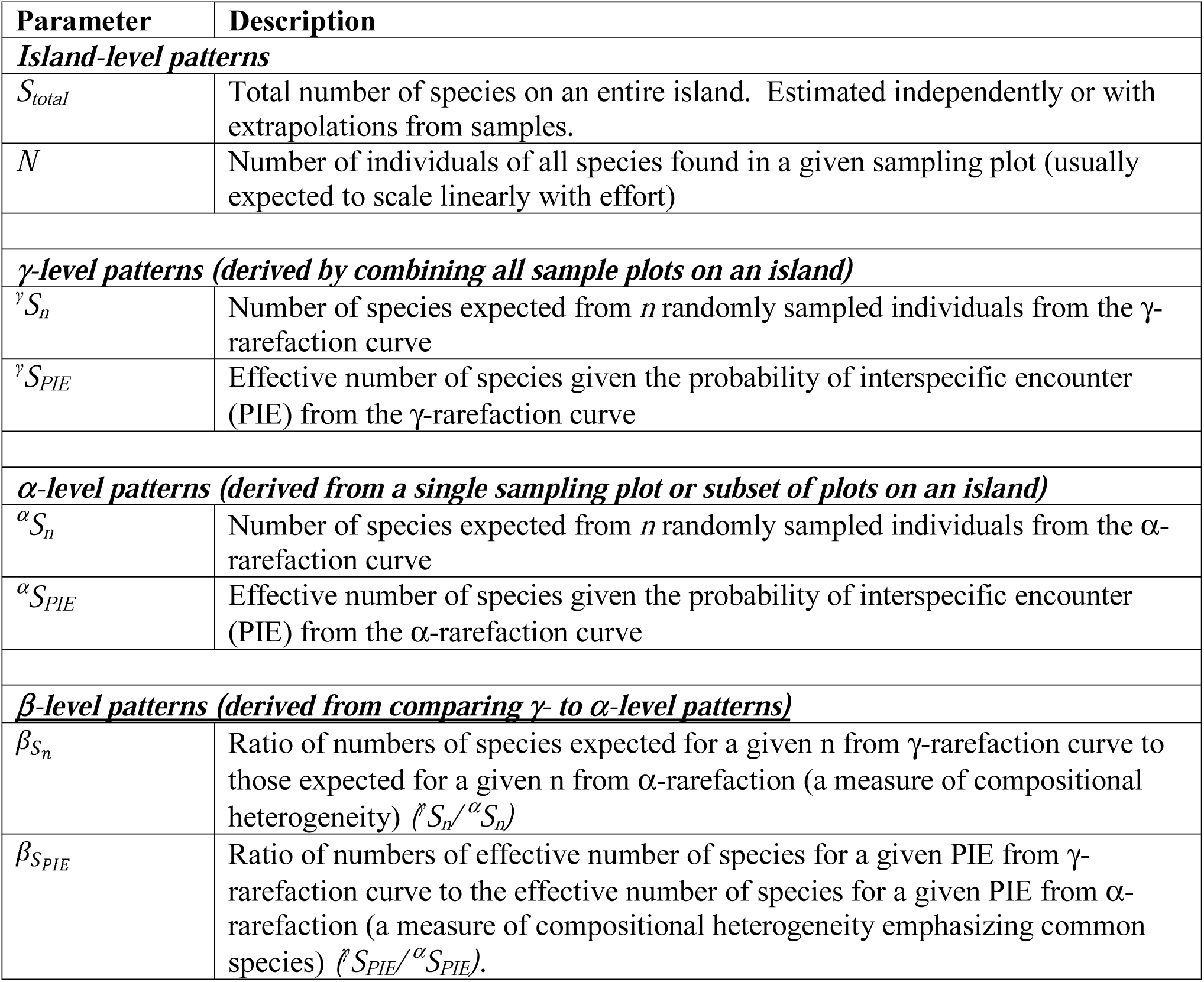
Parameters used to disentangle island species area relationship patterns

| Parameter | Description |
| --- | --- |
| ***Island-level patterns*** |  |
| $S_{total}$ | Total number of species on an entire island. Estimated independently or with extrapolations from samples. |
| $N$ | Number of individuals of all species found in a given sampling plot (usually expected to scale linearly with effort) |
| ***γ-level patterns (derived by combining all sample plots on an island)*** |  |
| ${}^{\gamma}S_n$ | Number of species expected from $n$ randomly sampled individuals from the γ-rarefaction curve |
| ${}^{\gamma}S_{PIE}$ | Effective number of species given the probability of interspecific encounter (PIE) from the γ-rarefaction curve |
| ***α-level patterns (derived from a single sampling plot or subset of plots on an island)*** |  |
| ${}^{\alpha}S_n$ | Number of species expected from $n$ randomly sampled individuals from the α-rarefaction curve |
| ${}^{\alpha}S_{PIE}$ | Effective number of species given the probability of interspecific encounter (PIE) from the α-rarefaction curve |
| ***β-level patterns (derived from comparing $\gamma$ to α-level patterns)*** |  |
| $\beta_{S_n}$ | Ratio of numbers of species expected for a given $n$ from γ-rarefaction curve to those expected for a given $n$ from α-rarefaction (a measure of compositional heterogeneity) (${}^{\gamma}S_n/{}^{\alpha}S_n$) |
| $\beta_{S_{PIE}}$ | Ratio of numbers of effective number of species for a given PIE from γ-rarefaction curve to the effective number of species for a given PIE from α-rarefaction (a measure of compositional heterogeneity emphasizing common species) (${}^{\gamma}S_{PIE}/{}^{\alpha}S_{PIE}$). |

**Figure 1.**
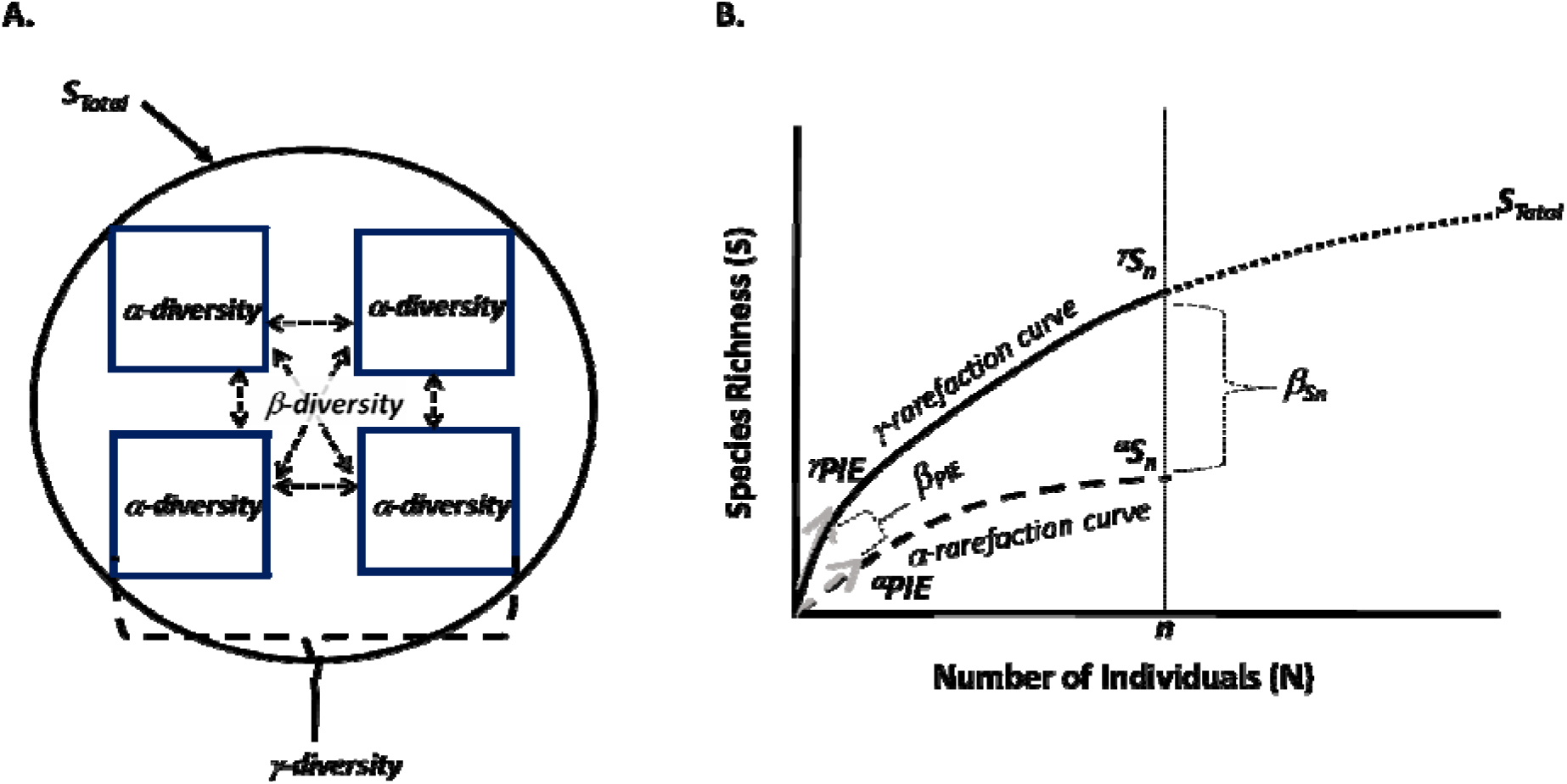
A. Overview of a sampling scheme appropriate for applying the analytical approach outlined in this paper. The circle represents a hypothetical island, and each of the four squares represents individual sampling plots from which α-diversity metrics can be derived. The addition of all of the individuals sampled in all of the plots allows the calculation of γ-diversity metrics, while the differences among the α-diversity plots is β-diversity. S_total_ represents the total number of species on the island, including those that were not observed in any of the sampled plots. B. Illustration of how these diversity indices can be visualized graphically from individual-based rarefaction curves that plot species richness (*S*) against the numbers of individuals (*N*) across scales. The γ-rarefaction curve (solid line) is derived by combining all individuals from all plots measured on a given island and randomizing individuals to generate the curve. From this curve, the dashed line allows us to visualize the total number of species on the island including those that were not sampled in any plot (*S*_*total*_). We can also visualize: (i) the numbers of species expected from a given *n*, γ*S*_*n*_ (where the vertical dashed line at *n* intersects the solid curve) (ii) the probability of interspecific encounter (PIE), which represents the slope at the base of the rarefaction curve, ^γ^*PIE* (solid grey arrow). The α-rarefaction curve (dashed line) is derived by randomizing individuals from a single plot, and similar parameters can be derived —^α^*S*_*n*_ (vertical dashed line intersects the dashed curve) and ^α^*PIE* (dashed grey arrow). The ratio between the γ- and α-rarefaction curves provides estimates of β-diversity that indicate the degree of intraspecific aggregation on the island. Note, in text, we advocate converting PIE values into effective numbers of species (*S*_*PIE*_), but only illustrate PIE in the figure, as it is not straightforward to illustrate *S*_*PIE*_ on these axes.

From the combination of all sampled plots within an island, one can generate a γ-rarefaction curve and several diversity parameters that can be derived from that information. We refer to the rarefied number of species expected from *n* randomly sampled individuals from the γ-rarefaction curve as ^γ^*S*_*n*_. Because the γ-rarefaction curve is generated by combining all sample plots on a given island and randomly choosing individuals, any spatial heterogeneity in species associations is broken. In addition to ^γ^*S*_*n*_, which weights common and rare species equally, we can also derive a measure which weights common species more heavily than rare species. Specifically, Hurlbert’s (1971) Probability of Interspecific Encounter (PIE) is a measure of evenness in the community and is equivalent to the slope of the rarefaction curve at its base, as illustrated by the gray arrows in Figure 1b (e.g., Gotelli and Graves 1996, Olszewski 2004). We use the bias-corrected version, 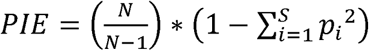, where N is the total number of individuals in the entire community, *S* is the total number of species in the community, and *p*_*i*_ is the proportion of each species *i*. For analyses, we convert PIE to an effective number of species (the number of species that would be observed if all of the species in a sample were equally abundant) (Jost 2006), which we call *S*_*PIE*_ *(=1/(1-PIE))*. PIE is the same as 1-Simpson’s diversity index, and when converted to an effective number of species, is part of the Hill continuum of diversity numbers that places more weight on common species (whereas richness places equal weight on common and rare species) (e.g., Hill 1973, Jost 2006). When *S*_*PIE*_ is calculated from the γ- rarefaction curve, we refer to the effective number of species as ^γ^*S*_*PIE*_. Note that only PIE, not *S*_*PIE*_, is illustrated in Figure 1b, because the forms of *S*_*PIE*_ are not readily illustrated in the individual-based rarefactions construct.

To discern whether any of the ISAR patterns emerge from within-island heterogeneity in species composition, we need to derive estimates of β-diversity. To do so, we can generate an α- rarefaction curve and estimate diversity parameters similar to those above, but at the local (within plot) scale. From this, we can compare the parameters from the γ-rarefaction curve which eliminates any plot-to-plot variation due to heterogeneity in species composition by randomizing across the plots, to the α-rarefaction curve calculated from individual plots (or a spatially defined subset of plots) which contains local information only (dashed line in Figure 1b). The degree to which the γ-rarefaction curve (which eliminates spatial heterogeneity) differs from the α-rarefaction curve (which keeps spatial heterogeneity) tells us how much local variation there is in species composition across sites, providing an index of β-diversity resulting from species aggregations (see Olszewski 2004, Chase et al. 2018, McGlinn et al. 2018). If the γ-and α-rarefaction curves are on top of each other, then we can conclude that there is no heterogeneity in the region. Alternatively, if the α-rarefaction curve is far below the γ- rarefaction curve, this implies that intraspecific aggregation has created compositional heterogeneity in the community. Two β-diversity parameters are informative in this context: 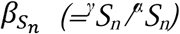 which indicates the influence of aggregation of all species, and 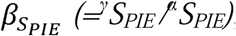, which indicates aggregations primarily by more common species (i.e., the effective number of unique communities; Tuomisto 2010).

In what follows, we discuss how this analytical methodology can be used to disentangle ISAR relationships where explicit sampling information from within and among islands is available. At the outset, it is important to note that in most of what follows, we focus exclusively on island systems where the primarily independent variable influencing species diversity is island size, with minimal variation in other diversity drivers. We focus on this because our goal is to elucidate and disentangle the ISAR, which describes a bivariate relationship between island size and species richness, and for which there remains much confusion and little synthesis. Nevertheless, as with all diversity studies, focusing on a single independent driver is a limiting case. In many island systems, islands vary in size as well as other drivers (e.g., productivity, isolation). Nevertheless, it is quite straightforward to extend the approach that we advocate below to include these complexities and still disentangle the influence of island size in the context of the ISAR. In such cases, one could simply use these other potential drivers as covariates with island size in an analysis focusing on the response variables we overview in Table 1 and Figure 1, using the same framework as described below. Or one could add more complexity by including these independent variables in a hierarchical model or structural equation model with the same response variables, which we discuss in more detail in the conclusions below (see e.g., Blowes et al. 2017, Chase et al. 2018 for similar analyses in a different context).

### Question 1: What is the shape of the overall ISAR? Parameter analyzed: Total number of species on an island (S_*total*_)

*S*_*total*_ is the most straightforward ISAR variable one can measure. The ideal way to estimate *S*_*total*_ is from independent information, such as exhaustive searching or checklists of species known to occur on a given island. However, because this information is often unavailable, *S*_*total*_ can be estimated via techniques for predicting the number of species in a given extent (e.g., Colwell and Coddington 1994, Harte et al. 2009, Chao and Jost 2012, Chao and Chiu 2014, Azaele et al. 2015). None of these approaches are perfect, and we are agnostic as to which approach is best for estimating *S*_*total*_ when complete species lists are not available. However, in our case studies below, we use the Chao (1984) non-parametric estimator to extrapolate the total number of species on a given island because it can be mathematically and conceptually linked to the rarefaction curves that we use (Colwell et al. 2012). However, this can only be viewed as a minimum expected *S*_*total*_, and will likely underestimate the true *S*_*total*_.

While *S*_*total*_ is the fundamental parameter of interest to calculate an ISAR, it alone provides little information as to the nature of its potential underlying mechanisms. This is because *S*_*total*_ is influenced by a number of underlying parameters, including the density of individuals, the relative abundances of species, and the intraspecific aggregation or spatial heterogeneity exhibited by species. Thus, to disentangle the factors underlying variation in *S*_*total*_, we need to look deeper into these underlying components, which we can do using the parameters overviewed in Table 1 and Figure 1b (see also Chase et al. 2018, McGlinn et al. 2018).

### Question 2: Does the ISAR result differ from what is expected from random sampling? Parameter Analyzed: Number of species expected from the γ-rarefaction curve (^γ^*S*_*n*_)

If patterns of the ISAR were generated simply by the random sampling hypothesis, we would expect that γ-rarefaction curves of small and large islands would fall right on top of each other (whereas the curve would go farther along the x-axis for the larger island, because more total N are present on larger islands) (Figure 2a). If the γ-rarefaction curves between smaller and larger islands differ, which we can quantify by comparing ^γ^*S*_*n*_ among islands (Figure 2b), then we can conclude that something other than random sampling influences the ISAR. This is essentially the same procedure as that described by the random placement approach (Coleman 1981, Coleman et al. 1982).

**Figure 2.**
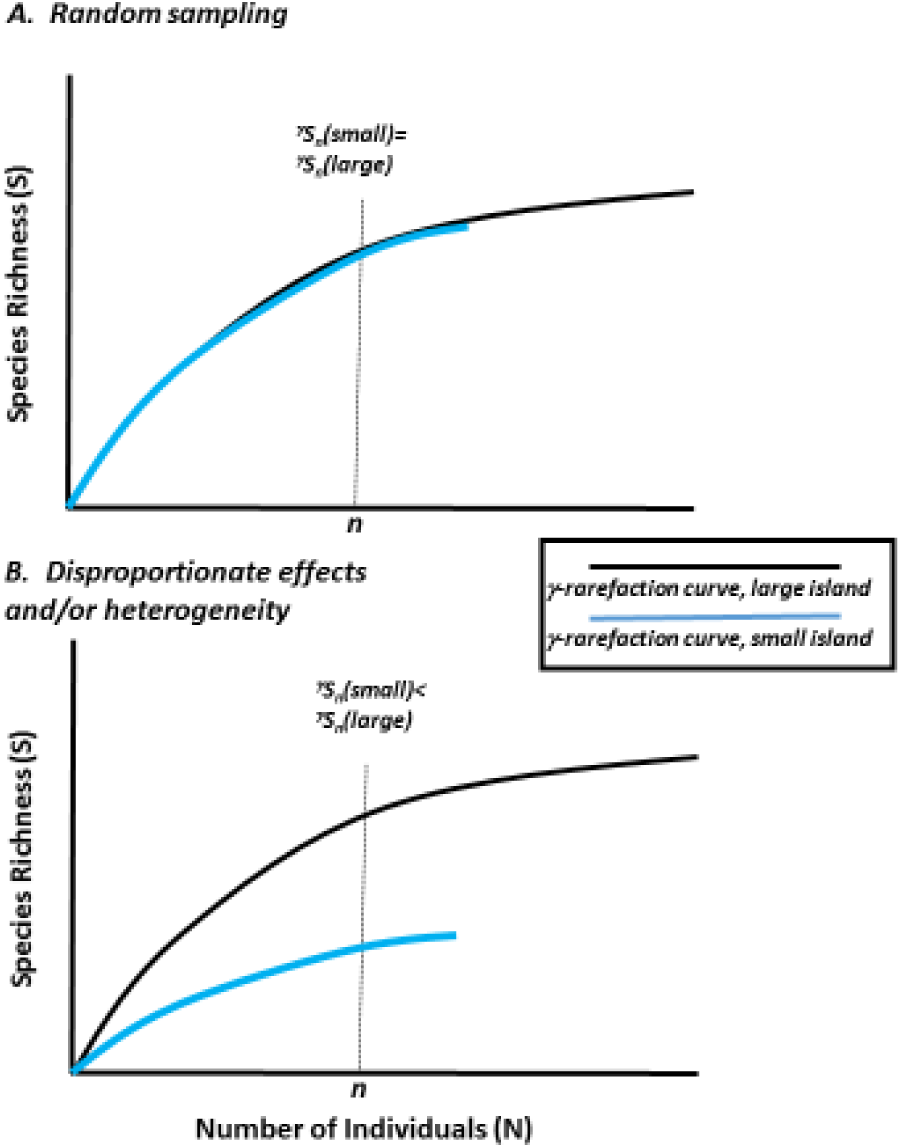
A. Hypothetical case where a large island has more species than a smaller island in total, but this entirely because of random sampling (the larger island has more total individuals). Note that the rarefaction curves for each island fall on top of each other, and the parameters derived from it, including ^γ^*S*_*n*_ and ^γ^*S*_*PIE*_ (not shown) are the same between larger and smaller islands. B. Hypothetical case where a large island has more species than a smaller island, and this results because both a sampling effect (the larger island has more *N*, and goes farther down the x-axis) as well as a disproportionate effect whereby ^γ^*S*_*n*_ is lower on the smaller than the larger island. ^γ^*S*_*PIE*_ in this case (not illustrated) is also smaller on the smaller island (because it has a shallower slope), but this need not be the case if only rarer species are affected.

If ^γ^*S*_*n*_ increases with increasing island area, this means that more species can persist for a given sampling effort on larger than smaller islands. We can go one step further in describing this pattern by examining how island size influences the relative commonness and rarity of species. If island area influences the γ-rarefaction curve via an overall decrease in evenness of both common and (as shown in Figure 2b), we would expect that both ^γ^*S*_*n*_ and ^γ^*S*_*PI*E_ would change. However, if only relatively rarer species are disproportionately influenced by island area (not shown in figure), we would expect that ^γ^S_n_ would increase with increasing island area, but there should be little to no effect on ^γ^*S*_*PIE*_. Importantly, the slope of the ^γ^*S*_*n*_ with island size very much depend on exactly which *n* is used in the calculations, with steeper slopes observed at higher *n*. This is similar to what was observed by Karger et al. (2014) on islands in Southeast Asia.

It is important to note that the hypotheses of increasing ^γ^*S*_*n*_ and/or ^γ^*S*_*PIE*_ with increasing island area, as illustrated in Figure 2b are not the only possibilities. Estimates of diversity from samples, such as ^γ^*S*_*n*_ and/or ^γ^*S*_*PIE*_, could certainly decrease with increasing island size. For example, on islands that result from habitat fragmentation and/or those that are surrounded by a relatively hospitable matrix, there are several mechanisms (e.g., habitat spillover) that can lead to higher levels of diversity (both in *S*_*total*_ as well as from samples [^γ^*S*_*n*_ and/or ^γ^*S*_*PIE*_]) in smaller relative to larger islands (e.g., Ewers and Diham 2006, Fahrig 2017).

Even if the numbers of species (and evenness) for a given sampling effort (^γ^*S*_*n*_ and/or ^γ^*S*_*PIE*_) declines, this can be outweighed by the random sampling effect, leading to an overall increasing ISAR even with decreasing components of diversity with increasing area. This emphasizes the fact that ISAR mechanisms are not mutually exclusive. That is, random sampling effects are likely always operating (as evidenced by the increase in species richness with increasing N along the rarefaction curve), even when disproportionate effects and/or heterogeneity also influence the ISAR pattern. As such, we can use rarefaction curves to examine whether random sampling is the only mechanism operating, as it would be if there is no influence of island size on ^γ^*S*_*n*_, and as a result, conclude that differential effects and/or heterogeneity are not operating. However, we cannot conversely say that random sampling is not operating if there is a relationship between ^γ^*S*_*n*_ and island size. This is because random sampling effects are always operating anytime there are fewer species on a given island than the total numbers of species in the regional species pool.

Finally, our discussion above implicitly assumed that island size changes the total number of individuals on an island via passive sampling, but not the density of individuals in a given sampled area. However, there are also reasons that island size can influence individual density. For example, if larger islands are more favorable for some reason, the total numbers of individuals would increase both because island size increases, as well as because the density in a given sampled area increases. Alternatively, smaller islands could contain more individuals for a given area (higher density) if there is high spillover from the matrix into smaller islands, or if larger islands have less favorable habitats. In such cases, comparisons of ^γ^*S*_*n*_ are still necessary to test the null hypothesis of whether the ISAR results from random sampling or not. However, when *N* varies with island size, it will also be useful to compare estimates of *S* at the scale of the sample rather than the number of individuals (i.e., sampled-based estimates sensu Gotelli and Colwell 2001, McGlinn et al. 2018) to determine how changes in *N* influence the ISAR.

### Question 3: Does the ISAR result from disproportionate effects or from heterogeneity Parameter analyzed: β-diversity as the difference between the γ-rarefaction curve and α- rarefaction curve

If there is a relationship between ^γ^S_n_ and/or ^γ^S_PIE_ and island area, we can conclude that there is something other than random sampling influencing the ISAR. With only the parameters from the γ-rarefaction curve, however, we cannot yet discern whether this is due to disproportionate effects that are equally distributed across the island, or whether these effects emerge because of heterogeneity in species composition across the island (i.e., different species and relative abundances in different parts of the island). To disentangle disproportionate effects from heterogeneity, we must look more closely into the variation in species abundances and composition within an island—that is, within-island β-diversity.

If 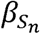 has no relationship with island size, then we can reject the heterogeneity hypothesis (Figure 3a; note, in the figure, we have illustrated that 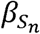 is 1, indicating there is no heterogeneity due to aggregation; however, this hypothesis would also be true if 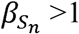, but does not significantly vary with island size). However, if 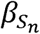 increases with island size, then we conclude that heterogeneity plays at least some role in the generation of the ISAR. If the ISAR is primarily driven by heterogeneity, we would expect there to be no relationship between ^α^*S*_*n*_ and island size, but a strong relationship with ^γ^*S*_*n*_, giving us a significant 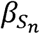 relationship with island size (Figure 3b). Such a pattern was observed by Sfenthourakis and Panitsa (2012) for plants on Greek islands in the Aegean Sea. In Figure 3b, we have illustrated a case where heterogeneity influences rare as well as common species, indicating an effect on both 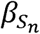 and 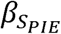 (not shown, but implied because the slope at the base of the curve [i.e., PIE] is influenced). However, it is also possible that heterogeneity can influence just the rarer, but not more common species, wherein we would expect an effect on 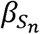, but not 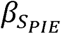 (not shown in Figure).

**Figure 3.**
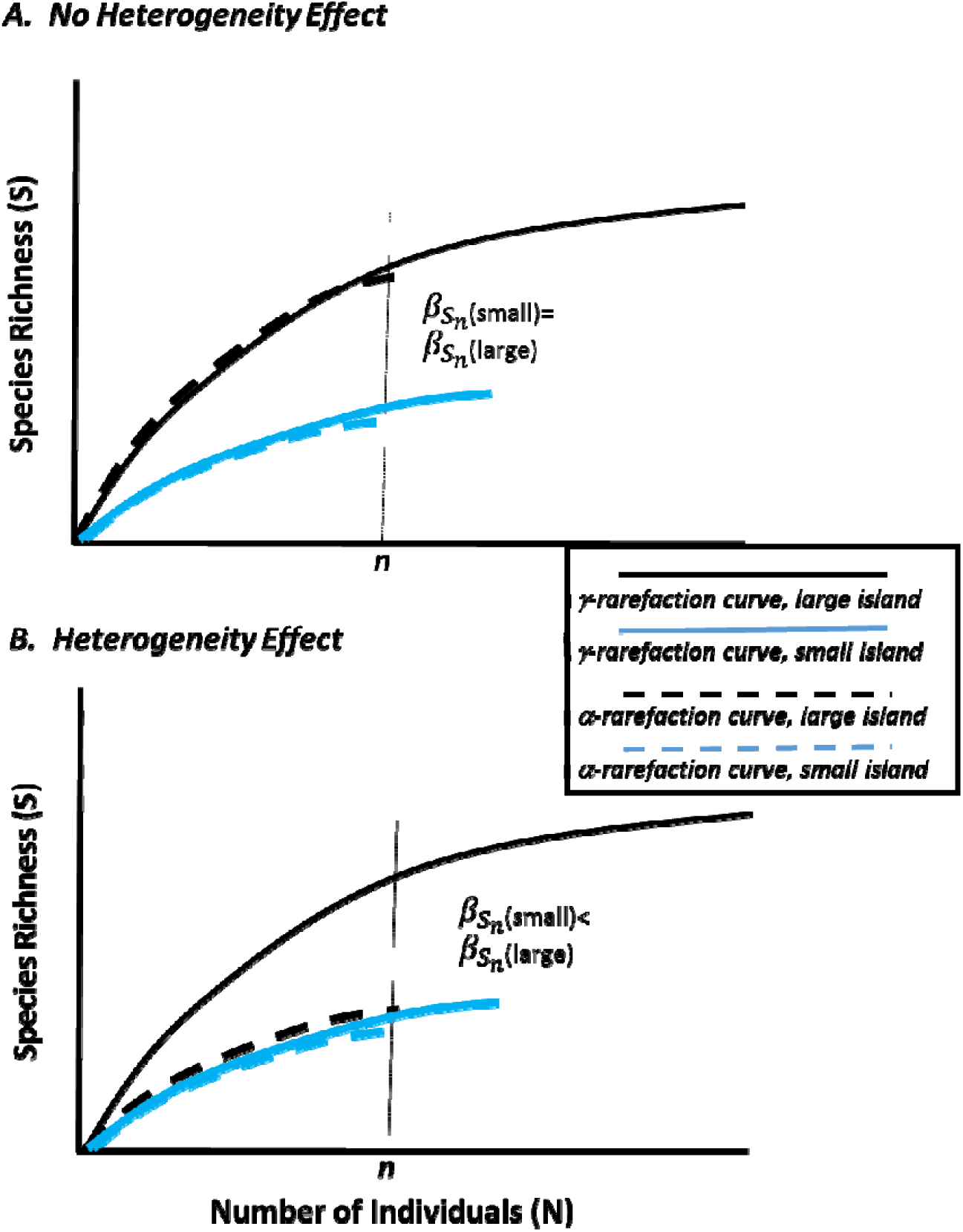
A. A hypothetical case where there is no heterogeneity in species composition within islands (the α- and γ-rarefaction curves completely overlap), such that =1. And this does not vary with island size. Note, that it is also possible that and/or >1, but we would conclude no heterogeneity effect underlying the ISAR if this is not influenced by island size B. A case where there is heterogeneity in species composition in the larger island (the α- and γ-rarefaction curves differ), but not the smaller. And thus, there is a positive relationship between compositional heterogeneity (and/or) island size. In this case, note that the α- rarefaction curves between the larger and smaller island overlap, and the island-effect is just observed at the γ-level, indicating the ISAR results solely from heterogeneity. This need not be the case, however, and other complexities can arise (see text).

It is quite possible that both disproportionate effects and heterogeneity occur simultaneously and in the same direction, in which case, we would expect a significant relationship between ^α^*S*_*n*_ and island size (indicating disproportionate effects) and stronger relationship between ^γ^*S*_*n*_ island size, giving a significant relationship between island size and 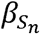 (not shown in Figure). On the other hand, disproportionate effects and heterogeneity mechanisms can act in opposition to one another. For example, the area-heterogeneity trade-off hypothesis assumes that as heterogeneity increases, the amount of area of each habitat type declines when total area is held constant (Kadmon and Allouche 2007, Allouche et al. 2012). Although perhaps not a common scenario (e.g., Hortal et al. 2009), if the types of habitats increase with island area, while the total amount of each habitat type declines, we might expect ^α^*S*_*n*_ and/or ^α^*S*_*PIE*_ to decline, while ^γ^*S*_*n*_ and/or ^γ^*S*_*PIE*_ can increase, remain unchanged or decrease, depending on the degree to which the heterogeneity effect is overcome by disproportionate effects (not shown).

Finally, if there is a significant relationship between 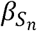 and/or 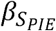 with island area, we can conclude that compositional heterogeneity likely underlies the ISAR, but we cannot infer whether this is due to habitat heterogeneity or dispersal limitation. To disentangle the relative importance of these mechanisms, it would be necessary to have additional information; for example, on the environmental conditions from different locations from within an island, and how species compositional heterogeneity was related to those conditions (see e.g., Leibold and Chase 2017 for an overview of approaches aimed at disentangling these).

### Caveat

Our approach, like all rarefaction-based analyses, assumes that sampling strategies can clearly identify and enumerate individuals of each species. Unfortunately, enumeration of individuals is much more difficult or impossible in certain kinds of communities (e.g., herbaceous plants, corals). Nevertheless, there are some ‘workaround’ solutions that can be used to apply the rarefaction techniques we have advocated to data where numbers of individuals are not available, but other measures of relative abundance are (e.g., percent cover or occupancy). For example, one can convert percentages of a species to individuals via a multiplier. In such a case, the meaning of PIE, S_n_ and aggregation measures change slightly, but can be calculated. Alternatively, one can collect presence-absence data on species in many quadrats within a locality. The presence of a species in a quadrat can be taken as a proportion and given the often strong correlation between abundance and occupancy (e.g., Gaston et al. 2000, Borregaard and Rahbek 2010), converted to an estimate of percent cover and converted as above. Again, while the interpretation of the parameters measured above cannot be taken literally, they provide a useful way to compare multiple diversity measures (at multiple scales) so that the framework we advocate can be applied.

## Case studies

Next, we illustrate how to use our analytical framework to test the ecological mechanisms underlying the ISAR with examples from three datasets representing different taxa and island settings. (1) Lizards sampled from several islands in the Andaman and Nicobar archipelago in the Indian Ocean (data from Surendran and Vasudevan 2015a,b); (2) Grasshoppers (Orthoptera) from Ozark glades, which are rocky outcrop prairies that represent island-like patches in a forested ‘sea’ (data from Ryberg and Chase 2007, Ryberg 2009); (3) plants from island-like habitat fragments of desert/Mediterranean scrub within an agriculture matrix (data from Giladi et al. 2011). For each case study, we present a brief overview of the system, results, and an interpretation of the results. We only used data from islands where multiple plots were censused; γ-measures included all of these plots, while α-measures were taken as the average among plots. Results are presented in Table 1 and Figure 4.

**Figure 4.**
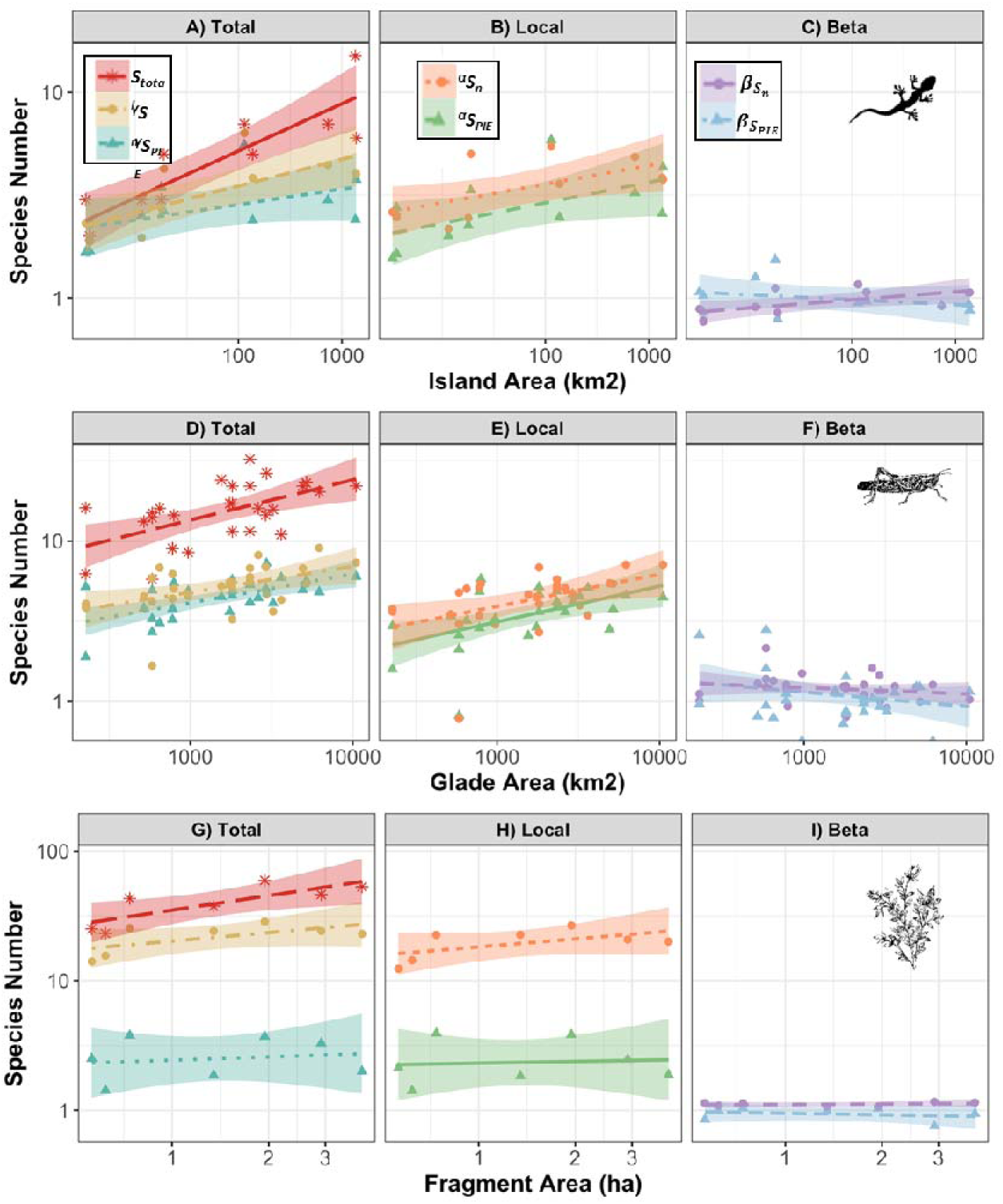
Log-log plots from the three case studies provided above. Each row represents results from a different case study; top row is for the lizards on the Andaman Islands; middle row is for the grasshoppers in Ozark glades; bottom row is for plants in Israeli fragments. Panels A,D,G represent parameters derived from the regional scale, including *S*_*total*_ (the number of species estimated on the total island), ^*γ*^*S*_*n*_ (the number of species expected for a minimum N measured across plots), and ^*γ*^*S*_*PIE*_ (the effective number of species given PIE across plots) (see text for explanation). Panels B, E, H represent parameters derived from the local scale, including ^α^*S*_*n*_ (the number of species expected for a minimum *n* measured in a single plot) and ^α^*S*_*PIE*_ (the effective number of species given PIE within a plot). Panels C, F, I represent parameters derived from comparing the local and regional scale (=β-diversity), including (the difference which represents heterogeneity in rare and common species), and (the difference which represents heterogeneity in common species). Coefficients and significance values are given in Table 1. Images are CC0 Creative Commons, with no attribution required.

All analyses were performed in R version 3.5.0 (R Core Team (2018). For each system, we did not have independent estimates of *S*_*total*_, and so we used extrapolation using iNEXT (Hsieh et al. 2018). We also used iNEXT to calculate rarefied richness values (*S*_*n*_). *S*_*PIE*_ was calculated using Vegan (Oksanen et al. 2017). Plots were generated using ggplot2 (Wickham 2016). Code and data for these analyses are available at https://github.com/Leana-Gooriah/ISAR_analysis

### Lizards on Oceanic Islands

The Andaman and Nicobar islands are a relatively pristine island archipelago in the Indian Ocean. A variety of taxa on these islands have been the subject of island biogeography studies, including ISAR studies (e.g., Davidar et al. 2001, 2002). Here, we used data from Surendran and Vasudevan (2015a,b) who intensively sampled lizards in 100 m^2^ quadrats from 11 islands that varied from 3.3 to 1375 km^2^ in area. We only used data from islands where two or more quadrats were censused.

As expected, we found a strong increase in our estimate of *S*_*total*_ as island size increased. We also found that ^γ^*S*_*n*_ increases significantly with island area, allowing us to reject the null hypothesis that the ISAR is driven by random sampling effects. However, the relationship between ^γ^*S*_*PIE*_and island area was not significant (Table 2, Figure 4a). A slightly different pattern emerged at the local scale (Figure 3b), with individual quadrats on larger islands having more species (^α^*S*_*n*_) that were less uneven in species composition (^α^*S*_*PIE*_) than on smaller islands. Because there were significant relationships between island size and both the γ-scale and α-scale measures, we can conclude that disproportionate effects played at least some role in driving the ISAR on these islands. Without additional information, we cannot say for certain exactly which spatial mechanisms are operating to allow more even communities, and more species co-occurring in local quadrats on larger compared to smaller islands. However, because 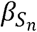 also increased with island size, this indicates that there was at least some influence of heterogeneity on the ISAR. This heterogeneity effect was only observed among the rarer species, but not the more common species, because there was no concomitant relationship between 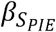 and island size. From other studies in these islands, we know that habitat heterogeneity does generally increase with island size (Davidar et al. 2001, 2002), and so suspect this relationship influenced heterogeneity in lizard composition from quadrat to quadrat more on larger relative to smaller islands.

**Table 2:** Linear regression coefficients and fits for each response in each case study. Coefficients are only given when the slope was significantly different from zero.

| System | Response | Intercept | Slope | $R^2$ | p-value |
| --- | --- | --- | --- | --- | --- |
| **Lizards on Oceanic Islands** | $S_{total}$ | 0.59 | 0.23 | 0.76 | **<0.0001** |
| | $\gamma S_n$ | 0.69 | 0.15 | 0.51 | **0.008** |
| | $\gamma S_{PIE}$ | - | - | - | 0.1 |
| | $\alpha S_n$ | 0.86 | 0.09 | 0.36 | **0.003** |
| | $\alpha S_{PIE}$ | 0.61 | 0.10 | 0.28 | **0.05** |
| | $\beta_{S_n}$ | -0.20 | 0.04 | 0.34 | **0.03** |
| | $\beta_{S_{PIE}}$ | - | - | - | 0.37 |
| **Grasshoppers**<br>**in Ozark**<br>**Glades** | $S_{Total}$ | 0.87 | 0.25 | 0.30 | **0.0001** |
| | $\gamma S_n$ | 0.45 | 0.16 | 0.19 | **0.01** |
| | $\gamma S_{PIE}$ | 0.18 | 0.18 | 0.35 | **0.0005** |
| | $\alpha S_n$ | -0.02 | 0.2 | 0.18 | **0.013** |
| | $\alpha S_{PIE}$ | -0.39 | 0.22 | 0.25 | **0.004** |
| | $\beta_{S_n}$ | - | - | - | 0.54 |
| | $\beta_{S_{PIE}}$ | - | - | - | 0.33 |
| **Plants in**<br>**fragmented**<br>**scrubland** | $S_{total}$ | 3.61 | 0.24 | 0.37 | **0.03** |
| | $\gamma S_n$ | - | - | - | 0.13 |
| | $\gamma S_{PIE}$ | - | - | - | 0.71 |
| | $\alpha S_n$ | - | - | - | 0.17 |
| | $\alpha S_{PIE}$ | - | - | - | 0.85 |
| | $\beta_{S_n}$ | - | - | - | 0.57 |
| | $\beta_{S_{PIE}}$ | - | - | - | 0.56 |

### Grasshoppers in Ozark Glades

Ozark glades are patchy island-like habitats within Midwestern forested ecosystems that contain xeric adapted herbaceous plant communities together with associated fauna (Ware 2002). Grasshoppers are diverse and abundant herbivores that are known to respond to local and spatial processes in these patchy ecosystems (e.g., Östman et al. 2007, Ryberg and Chase 2007). Here, we use data collected by Ryberg (2009) from area-standardized sweep sample transects (each sample represented 50 sweeps taken from a transect covering approximately 50 m^2^) taken from within glades without predatory lizards which varied from 0.02 to 1.05 ha (ranging from four transects on the smallest glade to 32 on the largest).

As with the lizards above, we find that S_total_ increases with island size, as does ^γ^S_n_. Furthermore, in this case, ^γ^S_PIE_ also increased with island size (Table 2, Figure 4d). A similar pattern is reflected at the local scale (Figure 4e). Thus, again, we can reject the null hypothesis that the ISAR emerges from random sampling, and instead we see a clear signal for disproportionate effects influencing both the number of species and their relative abundances. We suspect that one reason for this was because we only used glades that were relatively isolated from one another, and these grasshoppers do not readily disperse through the matrix. And thus, local processes likely outweighed any regional-level sampling effects. Unlike the lizards, however, we found no effect of glade size on β-diversity of grasshoppers between sweep samples within a glade (Figure 4F), suggesting that the ISAR did not likely result from increased levels of heterogeneity in larger glades, but rather from spatial processes associated with disproportionate effects. This is despite the fact that we know that heterogeneity in microhabitat types does increase with increasing glade size (Ryberg 2009). We suspect that the fact that these grasshoppers are rather mobile within glades was a reason that heterogeneity in species composition did not emerge among glades that varied in size. Unfortunately, however, without further information, we cannot say exactly what sorts of mechanisms allowed more species of grasshoppers to persist in larger glades than expected from random sampling.

### Plants in fragmented scrubland

Xeric scrub habitat in Israel was once quite extensive, but has been severely fragmented such that remnant habitats can be thought of as islands within a sea of agriculture (mostly wheat fields). These fragments have been the subject of intensive research on a number of organisms, including plants and several groups of animals (e.g., Yaacobi et al. 2007, Giladi et al. 2011, 2014, Gavish et al. 2012). Here, we used data from the Dvir region from the study by Giladi et al. (2011) on plants identified and enumerated in two or three 225 m^2^ quadrats within seven fragments varying from 0.56 to 3.90 ha.

Again, we found that S_total_ increased with fragment area, indicating a positive ISAR relationship. However, in this case, there were no significant relationships with ^γ^S_n_ or ^γ^S_PIE_ (Table 2, Figure 4g), any of the metrics from the α-rarefaction curve (Figure 4h), nor any of the β-scale metrics (Figure 4i). In this case, then, we are not able to reject the null hypothesis and instead conclude that the ISAR in these fragmented habitats is most consistent with the idea of random sampling. Even though we used different (and in our opinion, more robust) analytical tools, our results are qualitatively similar to those derived by the authors of the original study (Giladi et al. 2011). In this case, these results would indicate one of two general possibilities. First, it could be that these plants disperse well enough across the matrix that habitat size does not strongly influence local population dynamics. Second, it could be that local population dynamics do not depend on the numbers of individuals and types of species in local neighborhoods, at least during the time scale in which habitat fragmentation has taken place.

## Discussion and Conclusions

The island species-area relationship (ISAR)—depicting how the numbers of species increase with the size of the island or habitat patch—is one of the most well-known patterns in biogeography. Understanding the ISAR and the processes leading to it is not only important for basic ecological knowledge, but is also of critical importance for biodiversity conservation in the context of habitat loss and fragmentation. Despite this, the study of the ISAR continues to be difficult to synthesize, primarily because of the confusion about the confounding influence of sampling effects and spatial scale on the ISAR. For example, previous syntheses of the ISAR in natural and fragmentation contexts have focused on estimates of species richness at the entire island scale (e.g., Triantis et al. 2012, Matthews et al. 2016). Other syntheses, however, have confounded species richness measurements from multiple scales and contexts, making comparisons within and among studies difficult (e.g., Drakare et al. 2006, Smith et al. 2005, Fahrig 2017). As we have shown here, it is important to understand and report how species richness was sampled in order to interpret ISAR results. This is particularly true in the realm of conservation biology, where the influence of habitat loss and fragmentation on biodiversity is a critically important, but also controversial topic. In fact, a great deal of the controversy (e.g., Haddad et al. 2015, 2017, Hanski 2015, Fahrig 2013, 2017, Fletcher et al. 2018) is likely attributable to different investigators using different sampling procedures, different analyses, and different spatial scales for their comparisons, and thus comparing apples to oranges.

We are not alone in this call for a more careful consideration of sampling when measuring and interpreting ISARs (Hill et al. 1994, Schroeder et al. 2004, Yaacobi et al. 2007, Giladi et al. 2011, 2014, Sfenthourakis and Panitsa 2012, Karger et al. 2014). However, our approach, using metrics derived from γ- and α-rarefaction curves, provides an important advance over previous approaches by allowing one to more explicitly examine the influence of sampling and scale on the outcome. As our case studies illustrate, we can use this approach to explicitly disentangle the main hypotheses suspected to underlie the ISAR (random sampling, disproportionate effects and heterogeneity). For example, the case study on fragmentation in Israeli scrub habitats indicated that random sampling was primarily responsible for the ISAR. Interestingly, this result is similar to that found by Coleman et al. (1982) in their original use of this approach to another fragmented system; birds on islands within a flooded reservoir. Such results might be expected if species can readily use the matrix between habitat islands, or can easily disperse among habitats. Alternatively, in both the lizard and grasshopper systems, species are less likely to use the matrix and dispersal is likely lower, influencing the observation that disproportionate effects and heterogeneity influence the ISAR. Nevertheless, these are just a few case studies we analyzed where appropriate data were available. A more complete exploration of the generality of the patterns and potential mechanisms leading to the ISAR will require more thorough analyses of natural islands and patchy landscapes, as well as habitat islands that are created by habitat loss and fragmentation. Such analyses will allow us to achieve a more general synthesis of the patterns and possible processes creating ISARs in natural and fragmented island landscapes, but will also require more data (i.e., spatially explicit data of total and relative abundances of species as well as spatially explicit environmental data) than is typically analyzed in such studies.

There are clearly some extensions that can be made to the simple approach that we have overviewed. For example, when measuring ISARs in the real world, there are often many other mechanisms that can influence diversity patterns in addition to island size. For example, another important variable that influences diversity on islands is the isolation (distance) of those islands from others (e.g., MacArthur and Wilson’s 1967, Kreft et al. 2008). Habitat area can also influence trophic structure (e.g., larger islands may be more likely to have top predators), which in turn will feedback to influence the shapes of the rarefaction curves and patterns of diversity (e.g., Östman et al. 2007, Gravel et al. 2011). Likewise, in volcanic archipelagos, larger islands tend also to be younger and may have not had as much time for diversification as smaller/older islands, and this confounding factor can also greatly influence the shape of the ISAR (e.g., Whittaker et al. 2008,. Gillespie and Baldwin 2010). In addition, islands can vary in a number of other environmental and biological features, all of which can interact with island area. Fortunately, the metrics for which we have advocated which explicitly incorporate sampling theory and scale (see also Chase et al. 2018) can be analyzed in more complex models than the simple regressions that we have presented above. For example, hierarchical models can be applied to each of these metrics, analyzing the influence of island area along with a number of potential independent variables (see e.g., Blowes et al. 2017 for such analyses addressing a different set of questions). Likewise, structural equation models comparting patterns of ISARs along with several other covariables (e.g., Stiles and Scheiner 2010) can be applied to these metrics to disentangle area effects from other potential drivers.

Finally, despite its advantages, it is important to note that our approach is purely observational. As such, although it can provide deeper insights into the likely mechanisms that influence the ISAR than previous observational approaches, it cannot definitively discern process from these patterns. To more definitively test the primary ISAR mechanisms described here, we would need to go a step or two further. This could include, for example, observational studies that take advantage of variation, such as islands that varied semi-orthogonally in both area and heterogeneity (Nilsson et al. 1988, Ricklefs and Lovette 1999, Kallimanis et al. 2008, Hannus and Von Numers 2008, Stiles and Scheiner 2010), but also disentangling patterns of species richness in a more scale-explicit way as we have outlined here. Or it could include manipulative experiments that directly alter island size and/or heterogeneity (e.g., Simberloff 1976, Douglas and Lake 1994, Matias et al. 2010), or disrupt the processes occurring within islands (e.g., altering patterns of within-island dispersal and/or extinction).

## Acknowledgements

This work emerged from discussions among the co-authors in many contexts over many years, and was also improved by discussions with many other colleagues, including S. Blowes, T. Engel, P. Keil, S. Kroiss, D. McGlinn, B. McGill and N. Gotelli. Comments from R. Field, J. Hortal, S. Scheiner and an anonymous reviewer greatly helped us to improve the presentation. JMC, TMK, DC, LG, and FM were supported by the German Centre of Integrative Biodiversity Research (iDiv) Halle-Jena-Leipzig (funded by the German Research Foundation; FZT 118). The contribution of TMK and DC were also supported by the Helmholtz Association and by the Alexander von Humboldt Foundation. Ideas presented in this manuscript were inspired from work done as part of a grant supported by the U.S. National Science Foundation (DEB 0949984) to JMC and TMK.

## Author Contributions

JMC and TMK developed the initial conceptualization of the framework presented here, with significant input from WAR, MAS, FM, DC and LG at different stages. WAR provided the data for the grasshopper case study; FM provided the data for the fragmentation case study. LG wrote the code, with help from DC and FM, and did the analyses. JMC wrote the first draft of the manuscript, and all authors contributed significantly to revisions.

## Data and Code Accessibility

The code to run the analyses described here, as well as the data for the case studies, are freely available on https://github.com/Leana-Gooriah/ISAR_analysis.

